# Cyclical environments drive variation in life history strategies: a general theory of cyclical phenology

**DOI:** 10.1101/450387

**Authors:** John S. Park

**Author notes:** Corresponding Author*: John S. Park 1101 E. 57^th^ Street Zoology 400 Chicago, IL, 60637, USA. Statement of Authorship*: JSP conceived the idea, designed the study, performed modelling work and analyzed results, collected and analyzed field data, and wrote the manuscript.

## Abstract

Cycles, such as seasons or tides, characterize many systems in nature. Overwhelming evidence shows that climate change-driven alterations to environmental cycles—such as longer seasons— are associated with phenological shifts around the world, suggesting a deep link between environmental cycles and life cycles. However, general mechanisms of life history evolution in cyclical environments are still not well understood. Here I build a demographic framework and ask how life history strategies optimize fitness when the environment perturbs a structured population cyclically, and how strategies should change as cyclicality changes. I show that cycle periodicity alters optimality predictions of classic life history theory because repeated cycles have rippling selective consequences over time and generations. Notably, fitness landscapes that relate environmental cyclicality and life history optimality vary dramatically depending on which trade-offs govern a given species. The model tuned with known life history trade-offs in a marine intertidal copepod *T. californicus* successfully predicted the shape of life history variation across natural populations spanning a gradient of tidal periodicities. This framework shows how environmental cycles can drive life history variation—without complex assumptions of individual responses to cues such as temperature—thus expanding the range of life history diversity explained by theory and providing a basis for adaptive phenology.

## INTRODUCTION

Natural populations in all systems must survive environmental fluctuations. Biologists have long known that a particularly common and powerful mode of fluctuations in nature is cyclical, such as seasons. Species around the planet exhibit predictable and sensitive life history transitions that are tightly associated with seasonal cycles, also referred to as phenology. Environmental cycles in fact occur beyond just the timescale of seasons, such as daily, tidal, lunar, flood, fire and decadal oscillations, and life histories of species are often associated with cycles at these timescales as well [1–6]. Despite the ubiquity of cycles in nature, and clear empirical evidence of the importance of cycles for life histories, we lack a general theory of how life history evolution is shaped by cycles.

Over the last few decades perturbations to environmental cycles due to climate change have driven dramatic life history changes such as phenological timing in many species [7–15]. In fact, phenological shifts are widely regarded as the most conspicuous and rapid consequence of climate change across marine, freshwater, and terrestrial systems [14]. Notably, different species’ phenologies are shifting in different directions, creating phenological mismatches with profound consequences on ecosystem function and health [7,11,16–19]. Disparate case studies of shifts that typically invoke individual-level responses to environmental cues such as temperature may be limited in their potential to explain general evolutionary forces due to system-specific idiosyncrasies. On the trailing edge of rapidly accumulating empirical evidence of shifts, questions regarding general mechanisms of life history evolution in cyclical environments have emerged to the forefront of theoretical population biology, biodiversity, and climate change science [20–22].

A first step in understanding the mechanics of life history evolution in cyclical environments may be to conceptualize cycles as sequential arrivals of harsh conditions whose periodicity is not reciprocally affected by local ecological dynamics. An example is the arrival of winter in seasonal systems. A typical consequence of such cyclical events for a population is heightened mortality as well as some perturbation to population structure (e.g. seedling mortality in plants [23]). This consequence not only reduces population size at a given time, but also impacts the long-term trajectory and fitness of the population [24,25]. It follows that, if periodic disturbance is an inherent feature of a habitat, fitness is determined by how well a resident population survives repeated demographic perturbations at regular intervals.

Population ecologists have long been interested in demographic dynamics in variable environments, including cyclically variable environments [22,26–31]. Life history theorists, on the other hand, have classically focused on how time-invariant (i.e. constant) perturbations on age-, size- or stage-classes of populations, mediated by trade-offs between biological processes, shape life history strategies broadly [32–36]. For example, theory predicts that heightened juvenile mortality should induce the evolution of reduced reproductive effort. Such predictions have been widely tested empirically, and effects are often strong, rapid, and heritable [37–42]. So far, modern models of life history evolution that do incorporate time-variance in the environment have mainly focused on how optimality predictions are altered by stochasticity (i.e. randomly variable environments), which yield convenient analytical probabilistic conclusions [22,31,43–45]. What is not well understood is how life histories are generally shaped by non-random cycles, despite biological attention to fundamentally cyclical environments such as seasonal systems [22], and the fact that parametric changes to cycles such as season length are repeatedly associated with life history changes across systems.

Here I explore the general relationship between periodicity of cycles and evolutionarily optimal life history strategy. Proximate triggers of phenological expression, such as plastic response to temperature cues, mechanistically vary widely across species and habitats [20]. By taking a demographic life history theory approach agnostic to system-specific plastic responses, I address the ultimate selective force behind phenological traits and their shifts, given that phenology is fundamentally a study of how life cycle transitions are fit to environmental cycles.

I hypothesize that environmental cycles influence reproductive values of individuals and thus what life history strategy should be evolutionarily optimal in a given cyclical regime. Reproductive value is the expected contribution of an individual at a particular age or stage to the population through current and future reproduction, determined by biological trade-offs and survival through time [36,46]. Reproductive value is a central evaluation for fitness and evolution because it represents the aggregate consequence of trade-offs among many important life history traits [47]. Naturally, the realization of current and future reproduction must depend on current and future environmental conditions for survival experienced by individuals. Thus I expect that, under predictably cyclical environments that periodically incur harsh conditions for survival, the period length of cycles should have a tractable influence on which life history strategy should perform best in a given environment. I analyze this relationship by calculating which life history strategy in a population confers maximum long-term fitness in a given periodic regime, and then studying how optimal life history changes as periodicity changes. I explore how various trade-off assumptions impact these optimality curves to understand how different species in nature—whose life histories are in reality shaped by different sets of trade-offs—may be differentially affected by the same change in periodic regime.

Next I test my theoretical predictions in the copepod *Tigriopus californicus* (Copepoda: Harpacticoida), a crustacean found in rock pools in the supralittoral (upper tidal) zone along the North American Pacific coast. Populations are disturbed periodically by wave-wash at high tide, and experience population decline and heightened juvenile mortality periodically. Periodicity of disturbance varies among populations depending on regional tidal patterns and pool height on the shore. *T. californicus* provides an ideal system to study life history variation in cyclical systems across populations due to its short generation time and short disturbance cycles, the rare opportunity to sample from homogenized whole populations, and ease of quick sampling and trait measurements yielding large amounts of within-and across-population data. Across 19 natural populations of *T. californicus* in two regions of northern Washington I ask: do disturbance cycle periodicity and known trade-offs together predict life history variation across populations?

## METHODS

### Model construction

To uncover general predictions of evolutionarily optimal life history traits in cyclical environments, untied to species-specific idiosyncrasies such as plastic responses to meteorological cues, I describe a hypothetical population in two linked stages of broad applicability: juveniles and reproducing adults. I consider continuous-time demographic dynamics of the stage-structured population and impose stage-specific mortalities at given periodicities (full model description in electronic supplementary material, section 1).

First I express constant-environment dynamics as a system of ordinary differential equations

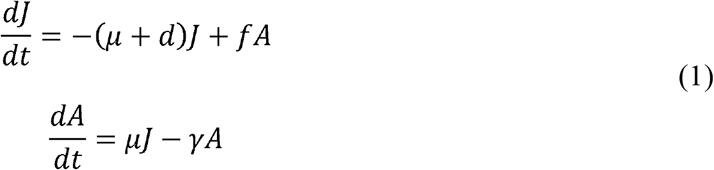

which can be expressed as matrix **M**:

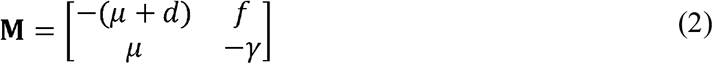

where J is juveniles, A is adults, *μ* is the rate at which juveniles mature into reproducing adults, *d* is background mortality of juveniles, *f* is the reproductive rate of adults, and *γ* is background mortality of adults. Then, via eigendecomposition of **M**, I express the solution at time *t* as:

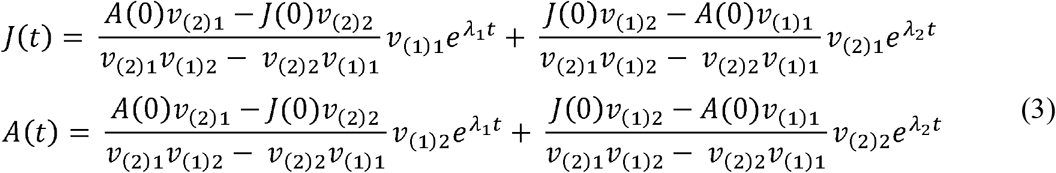

where v_(i)j_ is the j^th^ element of the i^th^ eigenvector corresponding to eigenvalue λ_i_ of **M**. This solution describes simple structured population dynamics in an undisturbed environment, but by eigendecomposing the system I isolate the time parameter *t* which will eventually allow me to study demographic dynamics as a direct function of period length between disturbances. To make the solutions explicit with respect to disturbance cycle period T, I let *t* = T, and at time T multiply the structure by S_J_ and S_A_ to impose juvenile- and adult-specific mortality associated with disturbance. The combined system can be expressed as the matrix **P** (S10):

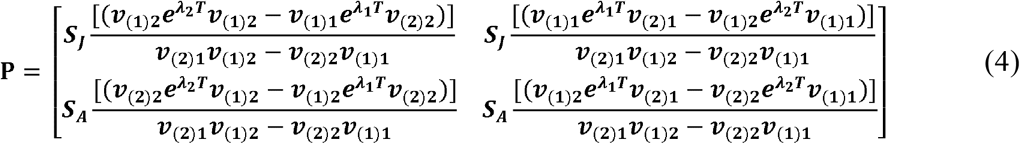

Matrix-multiplying initial abundances by **P** would thus give stage structure after existing in a constant environment for time T and experiencing a disturbance event that incurs stage-specific mortalities. More interestingly, I use this framework to ask: what are the consequences of different combinations of life history traits on the fitness of a population given that it resides in disturbance regime T?

### Fitness

Given the general framework of cyclically perturbed stage-structured population dynamics, I ask how the predicted fitness of the population is influenced by the periodicity of environmental cycles. The dominant eigenvalue (λ) of a population transition matrix is a widely used measure λ of relative fitness because it represents how well the population will perform in the long run compared to other hypothetical populations with different life history strategies [25,36]. This metric, equivalent to ‘*r*’ in demography and life history theory, does not capture consequences of short-term transient dynamics [48,49], but has been useful for drawing broad life history evolution predictions and conceptualizing relative fitness that match well with empirical observations [24,25,36]. In stochastic environments fluctuations in instantaneous growth rates may lead λ to give inaccurate evolutionary predictions. In systems that can be modelled by λ periodic switching between environments, however, eigenvalues and eigenvectors of the matrix product of constituent matrices describing the different environmental states can be used for demographic and life history analyses in exactly the same way as they are used in time-invariant theory [25,50]. My matrix **P** is equivalent to periodic models since the system switches between an undisturbed phase and disturbance, and the switching periodicity and population matrix elements do not fluctuate randomly (see electronic supplementary material, Fig. S1 for simulation results). Thus, here I use the dominant eigenvalue of **P** (hereafter referred to as λ_P_) as the measure of relative fitness to compare the theoretical performance of life history strategies in a periodically time-variant framework, and characterize general selective pressures on life history strategies as a function of cycle periodicity.

### Life history trade-offs

Life history evolution is a matter of optimization because limited resources must be allocated into various biological processes such as survival and reproduction involving trade-offs [36,51]. The exact shapes of trade-off functions in organisms are famously difficult to measure, let alone justify in model assumptions [51,52]. Here I take a conservative approach and assume simple linear trade-offs to investigate general patterns in optimality as a function of the environment without making more complex physiological assumptions. To express a trade-off between any two traits in the construction of a fitness landscape, I computationally set the vector of the range of values of one trait in decreasing order as the other increases, imposing a negative slope between the two traits. When two traits do not trade off, one of the traits remains at the mean of its range as the other varies through its own range. I varied the combinatory inclusions of trade-offs among the four key parameters to create model variants and investigate their relative fit to the data.

### Fitness landscapes and optimal life history strategies

All realizations of **P**—and thus the construction of fitness landscapes—must be constrained within the space of the interacting life history parameters, *μ, d, f*, and *γ*. In this presentation I constrained the space with known *T. californicus* life history ranges and trade-offs to demonstrate one example of the usage of this framework, but constraints can be set flexibly to represent any given species (see electronic supplementary material, section 2.5 for descriptions and citations for parameterization).

Using λ_P_ I construct fitness landscapes for *μ* and *f* simultaneously for each model. Here I focus on *μ* and *f* because they are life history traits for which I can collect large amounts of paired data in *T. californicus*, but it should be noted that fitness landscapes can be created for any life history trait in the original system of differential equations. For each landscape, I scan across the range of *μ* or *f* for a given value of T, while varying all other traits according to trade-off relationships included in the given model. Therefore I construct a vertical gradient of relative λ_P_ per T. To construct a landscape, I calculate gradients of relative λ_P_ across the horizontal axis of T. The optimal trait per T is the trait that maximizes λ_P_ per T. Finally, to get the curve of optimal trait values across the axis of T I track values associated with maximum λ_P_ across T.

### Empirical investigation in *Tigriopus californicus*

*Tigriopus californicus* is a copepod found widely along the North American Pacific coast (see electronic supplementary material, section 2.1 for detailed description of natural history). Dense populations reside in rock pools above the intertidal zone at varying heights [53–55], which accordingly experience tide cycle disturbance at varying periodicities. When tide levels cyclically reach pool heights and waves wash through pools, *T. californicus* cling onto the rocky benthos in order to prevent being flushed down to open water or to the lower intertidal zone [55]. If they are washed down, predators that do not occur in *T. californicus* pools feed on them quickly, and re-colonization of *T. californicus* into the pools appear to be low [55,56]. Despite clinging, tidal disturbance was shown to always decrease population size, and in particular, incur heightened juvenile mortality (electronic supplementary material, Fig. S4).

I sampled 19 isolated populations across two sites in northern Washington, USA (Neah Bay, Friday Harbor) in order to capture a wide gradient of disturbance periodicities (see electronic supplementary material, sections 2.2-2.4 for detailed description of data collection). I quantified the periodicity of tidal disturbance in each pool via timeseries analysis of pool temperature data over 4 months at 5-minute intervals, taking abnormal drops in temperature as signals of wave flush (see electronic supplementary material, section 2.2). I siphoned entire isolated populations out of rock pools, and subsampled individuals after homogenizing them, to get representative population samples. I reared 30 mating pairs captured from each population in common garden settings. In these lines I measured rate of maturity (*μ* in the model) and rate of reproduction (*f* in the model) (see electronic supplementary material, section 2.4 for detailed description of trait measurements).

### Likelihood and model fitting

I calculated the log-likelihoods of the optimality curves of the two focal life history traits *μ* and *f* produced by each model variant given the variance and covariance of the *μ* and *f* data. Each model is a different trade-off model (electronic supplementary material, Fig. S2, Table S2). Every model has the same number of estimated parameters because they only differ in how the parameters trade off in the construction of the fitness landscapes, which is included computationally by aligning parameter range sequences in reverse order. Therefore model selection criteria that penalize number of parameters such as AIC were not used. Each model produces optimality curves (dominant eigenvalue of matrix **P** across gradient of disturbance period T) of *μ* and *f* given trade-off relationships. I searched for the maximum log-likelihood of each model given *μ* and *f* data simultaneously within the space of S_A_ ≥ S_J_ and compared maximum log-likelihoods of the 13 model variants.

## RESULTS

### Cycle periodicity alters optimal life history predictions

Classic life history theory balances costs and benefits of key biological investments such as development, reproduction, and survival to predict fitness profiles of life history traits [36,57,58]. Here I incorporated these classic balance considerations but imposed cyclical perturbations to population structure and asked if the fitness predictions change as a function of environmental cycle periodicity. Using this framework, I analyzed the role of cost (slope of trade-off, Fig. 1A) on the fitness profile of a life history trait (maturation rate) in two scenarios: one in which period length is long enough (e.g. to fit more than 10 generations in a period) that the effect of discrete cycles on the evolution of life history rates should be small (Fig. 1B), and another in which period length is at a similar timescale to generation time (Fig. 1C). The former approaches classic formulations of optimal life history predictions based on trade-offs alone [57]. The latter shows that external periodic perturbations significantly change optimality predictions. In the latter scenario all trade-off cost assumptions predict higher optimal values of maturation rate compared to the former. The shape of fitness profiles is also flatter in the latter scenario, which may suggest weaker selection or that larger variance of maturation rate can be maintained within a population under shorter disturbance cycles. Lastly, the relationship between trade-off cost and optimality is reversed between the two scenarios: the lowest cost case produces the lowest optimal maturation rate under long periods but the highest optimum under short periods, and vice versa. These results show that the periodicity with which harsh environmental conditions arrive and affect survival modifies the expected reproductive value of individuals, and significantly alters relative fitness of strategies with which individuals invest biological resources into life history traits. This effect can reshape classic predictions of optimal life history that are solely based on internal trade-offs.

**Figure 1.**
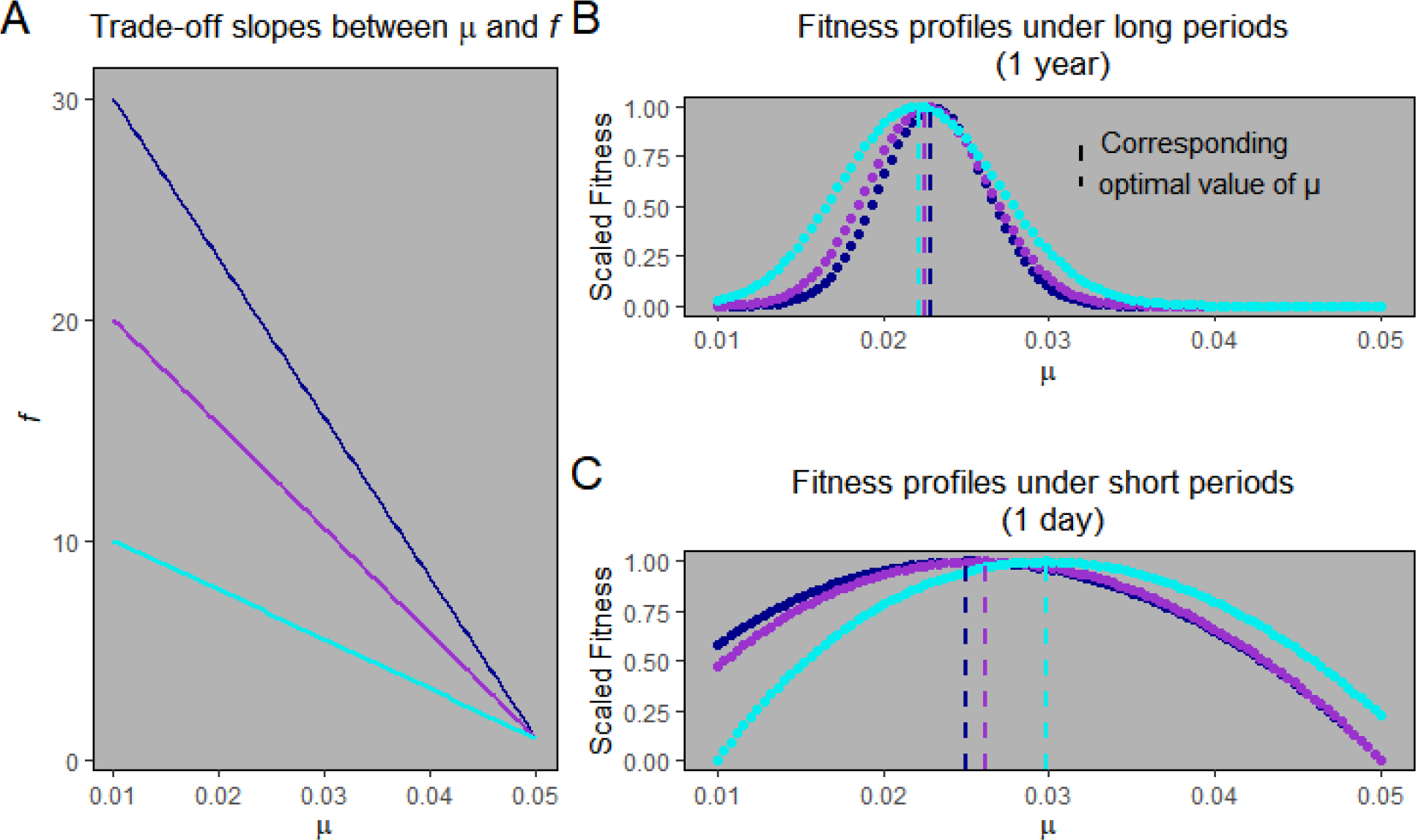
*Three hypothetical cost functions between* μ—*rate at which juveniles mature into reproducing adults—and* f— *adult fecundity—are analyzed while setting linear trade-offs between* μ *and* f, *and between those two traits and their respective stage-specific background survival rates (*d *and* γ*). Stage-specific survival terms associated with cyclical disturbance are set at S_A_ = 0.9 & S_J_ = 0.6. Colors of cost functions in A correspond to colors of fitness profiles of* μ *in B and C. Dashed lines in B and C show peaks of fitness profiles which correspond to optimal values of* μ*. Periodicity of cyclical perturbation to population structure is set to be much greater than generation time in B (T=365), and at a relevant time scale (<generation time) in C (T=1). Under short periods (C), all cost functions produce higher optimal* μ *values, wider fitness profiles, and an exactly reversed relationship between cost and optimality compared to long periods (B).*

### Periodicity and trade-offs interact to produce diverse life histories

Optimal life history varies nonlinearly as a function of disturbance cycle period, even with assumptions of simple linear trade-offs between traits (Fig. 2). This nonlinearity implies that changes in the evolutionary optimum of a life history trait can be of very different magnitudes even with the same magnitude change in periodicity, depending on the initial period length.

**Figure 2.**
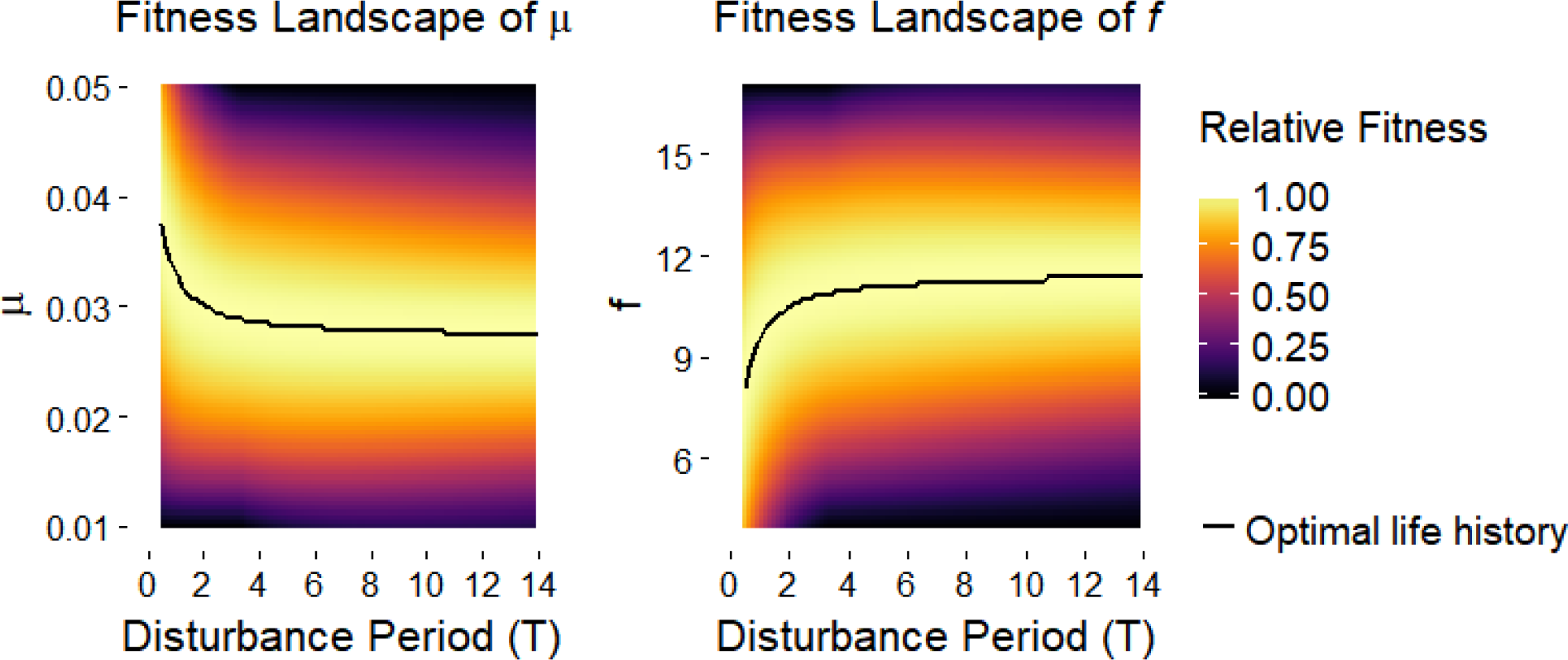
*Example fitness landscapes of two focal life history traits*, μ *(rate of maturity) and* f *(reproductive rate) which assumes lower juvenile survival with each disturbance event (S_A_ = 0.9, S_J_ = 0.6), and trade-offs between* μ *and* f*, between* μ *and* d*, and between* f *and* γ. *Heat shows normalized fitness of a life history strategy compared to all other strategies in a disturbance regime (T). Curves track the optimal (maximum fitness) life history trait across T*.

Shapes of optimality curves (optimal traits vs. period) can vary dramatically depending on which life history trade-offs are included (electronic supplementary material, Fig. S2). For example, when maturation trades off with background juvenile survival and fecundity (Fig. S2G), optimal maturation rate is expected to decrease and optimal fecundity is expected to increase as period length increases; on the other hand, if maturation trades off with background adult survival and fecundity instead (Fig. S2I), directions of expected trends in both optimal traits as period length changes are the opposite of the former case. Similarly, when maturation trades off with background adult survival, optimal maturation rate and fecundity are both expected to increase with period length (Fig. S2C), but both are expected to decline with period length if background juvenile survival trades off with background adult survival (Fig. S2E) or if fecundity trades off with background adult survival (Fig. S2F); on the other hand, both optimal maturation and fecundity are insensitive to change in period length if maturation trades off with background juvenile survival (Fig. S2A). Collectively, this broad range of cases examined demonstrates that the way in which external environmental cycles determine what combination of life history traits is evolutionarily optimal depends heavily on how traits trade off of one another internally. In the next section I show that the model that includes known trade-offs in *T. californicus* has the highest likelihood given *T. californicus*-specific life history data; but it is important to note that no one model is necessarily better than another in a general sense because different species in nature will have different levels of complexity and rank order of trade-offs between life history traits [57,51,59].

A long-accepted tenet in life history evolution theory is that the mean and variance of population structure perturbations shape life history variation [24,34,35,39]. Results here show that the temporal nature of such perturbations, such as the period length of environmental cycles, should interact strongly with general life history trade-off architectures in determining evolutionarily optimal traits. Trade-off patterns realistically vary widely among species due to variations in physiological and developmental mechanisms [51]. Periodicity of environmental cycles, such as growing season length across latitudes, varies widely across habitats as well. The interactive effect of environmental cycles and life history trade-offs is a previously unexplored cause of variability in optimal life history which can drive diverse strategies in nature.

### *Tigriopus* trade-offs predict life history variation across a periodicity gradient

Temperature time series analyses confirmed that there is a broad range of disturbance cycle periodicities across *T. californicus* pools across the two regions (electronic supplementary material, section 2.1; Fig. S4A, B; Table S1). These sampled pools provided a gradient of periodic regimes against which I tested optimal life history predictions. Daily temperature regimes, which may contribute to life history differences [60,61], were not significantly different among pools of varying periodicity regimes across the two regions (Fig. S5). Disturbance always caused higher juvenile mortality than adult mortality in subsampled disturbance events, with mean juvenile mortality of 41% and mean adult mortality of 6% (Fig. S4C).

Life history traits shift as disturbance period changes across *T. californicus* populations (Fig. 3), mirroring the shape predicted by the model (Fig. 2). The best model (likelihood maximizing when *μ* and *f* are fit simultaneously, represented by Fig. 2) was the one that assumed trade-offs between maturation rate and fecundity, between maturation and juvenile survival, and between fecundity and adult survival, consistent with known trade-offs in *T. californicus* (electronic supplementary material, section 2.1). Raw data collected for *μ* and *f* per maternal line in my populations also support a general negative relationship between *μ* and *f* (electronic supplementary material, Fig. S6). Finally, model variants with double or tertiary trade-off assumptions generally fit better than ones with only single trade-offs (see electronic supplementary material, Fig. S2 and Table S2 for the full list of models). These comparisons among model variants suggest that multidimensional trade-off relationships—which are typically avoided in empirical measurements or model assumptions of life history evolution [51,52] but gaining some attention [62,63]—may actually be important in predicting life history optimization in cyclical environments because trade-off consequences change as a function of cycle period.

**Figure 3.**
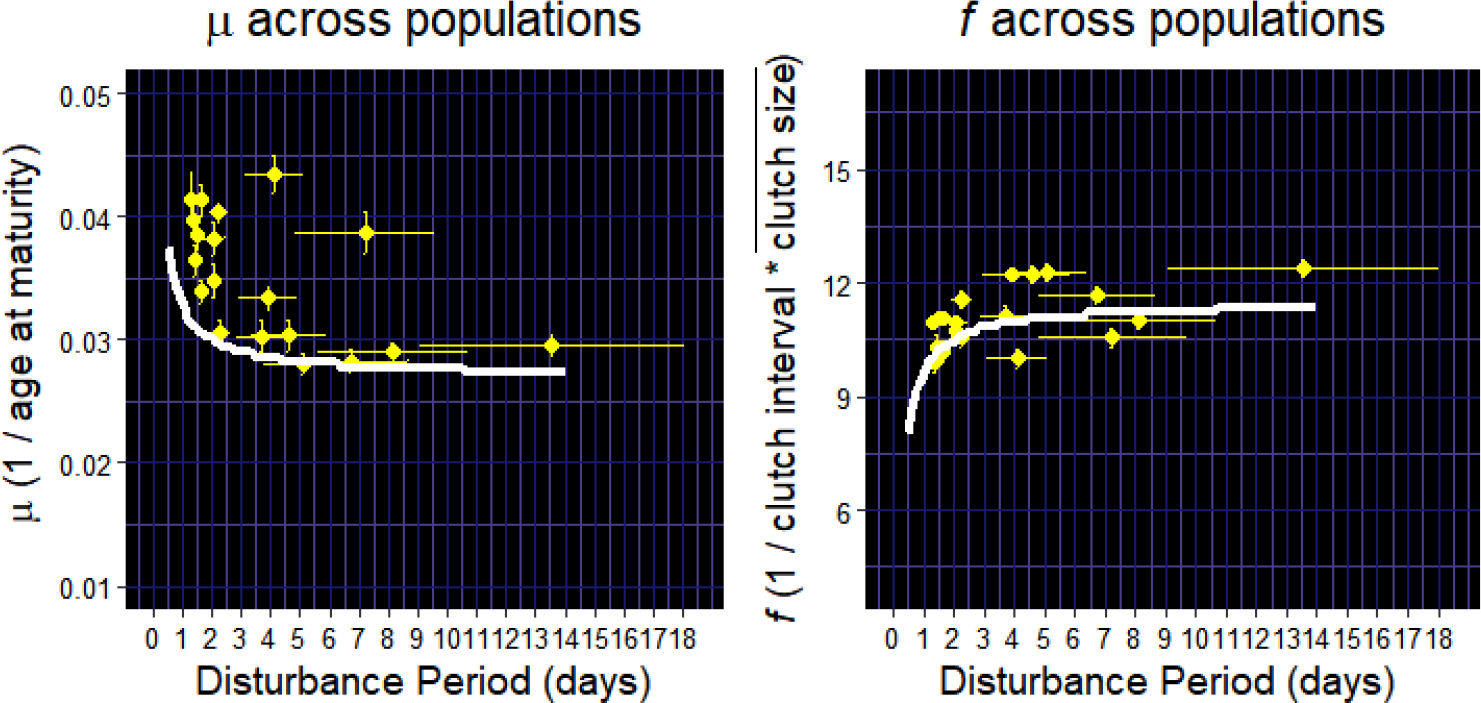
*Mean (±se) values of the two focal life history traits* μ *and* f *across 19* T. californicus *populations, against mean (±se) disturbance period determined by timeseries analysis of wave disturbance signals in each pool. Curves are optimal life history functions across periodicity (T) fit simultaneously to* μ *and* f.

## DISCUSSION

Ecologists have long assumed that environmental cycles are important for life cycle-related traits. But growing knowledge of phenological shifts has generated confusion regarding how environmental cycles shape life history strategies and thus transition rates of life cycle phases. One unresolved paradox in phenology is that various species in the same community (e.g. those in different trophic levels) undergoing the same change in seasonal cycles often exhibit phenological shifts in opposite directions. Here my results suggest that an interaction between environmental cycles and general biological trade-off relationships among fitness-related traits might contribute to life history and phenological divergence: the long-term consequence of trade-offs, a fundamental driver of life history strategy, is modified by environmental cycle periodicity, and significantly alters traditional predictions of optimal life history that are based on assumptions of population structure perturbations and trade-offs alone. A version of the model tuned with known *T. californicus* trade-offs successfully predicted the shape of life history variation across natural periodicity regimes, demonstrating the power of this interactive effect.

A fundamental question in ecology and evolution is why life histories are so diverse in nature. Divergent trends in phenological shifts among species in fact offer a current, global opportunity to study the production of life history diversity. Here I show that the interaction between environmental cycles and life history trade-offs is a simple mechanism that can account for large variations in life histories. First, due to the non-linear relationships between cycle period and optimal traits, the same magnitude of period change can induce different magnitudes of life history evolution between two populations of a species that are in different cyclical regimes (Fig. 2). Second, different trade-offs produce varying shapes of optimality curves (electronic supplementary material, Fig. S2), and thus the same change in period can induce an increase, decrease, or no change in a life history trait for different species in the same system depending on what trade-offs are biologically important for those species. Environmental cycle periodicity is diverse across systems (such as growing season lengths across a latitudinal gradient), and trade-off architectures among populations and species vary widely due to physiological constraints, environmental conditions, and reaction norms [51]. Combined, cycles and trade-offs can produce a wide array of predicted life history strategies. Testing this mechanism in species that are controlled by different trade-offs, either across populations in different cyclical regimes or within a single population through time in a habitat undergoing a change in cycle periodicity—for instance due to climate change—will provide fruitful avenues for further exploring this perspective.

### Stochasticity, ESS models, and gene flow

Cycles in nature, of course, are not perfectly periodic. The present study focuses on the consideration of period, or interval length between autocorrelated events. The mechanistic influence of fundamentally cyclical environments on life history evolution is noticeably understudied compared to probabilistic expectations in stochastic environments [22], even though regular cycles on various time scales are common in nature. Periodic models can be used to address a real aspect of nature that is difficult or impossible to address explicitly with stochastic models: cyclicality. Here, I take advantage of the fact that periodic models allow the use of matrix properties such as the dominant eigenvalue to infer relative fitness within a fluctuating system [25,50] and analyze conditions for optimization. By doing so I uncover a novel mechanistic relationship between cyclicality and life history evolution. However, cyclicality and stochasticity are both important aspects of nature. For instance, stochastic fluctuations in instantaneous population growth rate can significantly modify evolutionary trajectories predicted by time-invariant or periodic theoretical assumptions [48,49,64] Studying the relative influences of periodicity and stochasticity on optimal strategy, and on how quickly a population evolves to its predicted optimal strategy, are the obvious next steps that will add more richness to the perspective offered here.

Optimality curves in my model framework represent variations in evolutionary stable strategies (ESS) because I take the long-run growth rate of populations (dominant eigenvalue of **P**) as the measure of fitness as is commonly done in demography and life history theory. ESS models are useful for the purpose of predicting general directions of selection over a long term. ESS models take a non-genetic perspective on broad selective forces, although a genetic justification for optimization of a quantitative trait is given by the fact that a mutation can invade the population if it confers a higher *r* on its carriers [24]. Optimization models and quantitative genetics models are approximately equal for constrained multivariate systems [65]. Nonetheless, results found here are inconclusive with respect to what a population’s evolutionary trajectory from one optimum to another should look like in an environment undergoing change in cycle periodicity. Antagonistic selection on correlated traits imposed by different environmental variables associated with seasonal fluctuations, such as photoperiod and temperature, might cause deviations from ESS predictions. Evolutionary trajectories could be altered if bottlenecks are created by a sequence of disturbances and constrain the standing genetic variation subject to selection. In *T. californicus*, selection on optimal life histories may be obscured if high gene flow among nearby populations exists due to wave transport. However, colonization rates and genetic exchange have been repeatedly observed to be low in this system [66–68], and demographic dynamics given high mortality rates caused by tidal disturbance likely overwhelm population genetic dynamics on the time scale of tide cycles. In this study I deliberately chose populations that were deemed to be well isolated given field observations. But the level of gene flow may vary depending on locality due to habitat characteristics, and may contribute to some of the variance within populations and deviations of population means from ESS predictions. Nonetheless, my model fitting results suggest that ESS assumptions predict *T. californicus* life histories reasonably well given a population’s periodic regime.

### Trade-off functions

Trade-offs between traits can be nonlinear, and multidimensional architectures of trade-offs can be extremely difficult to measure [51,59,52]. Here I have taken the conservative approach of assuming linear trade-offs among modeled life history variables, which biologically equate to strictly substitutable energetic currencies divvied between different traits, to focus on the demonstration that consequent optimality curves across periodicity are nonlinear, and that a diverse set of optimality curves can be produced with different trade-offs. The simple linear assumption still performs well, at least with *T. californicus* life history data from my sample populations. However to test this framework further in different species, different functions can and should be used if the relationship between two traits is known to be nonlinear.

### Links to evolution of seasonal phenologies

In seasonal environments cyclical arrival of harsh meteorological conditions (e.g. winter) can incur large demographic perturbations and thus strongly influence population dynamics [69,70]. Here I show that if periodic arrivals of disturbance incur significant demographic perturbations, individuals and their lineages that have life history strategies that are non-optimal in the context of their environment’s cyclicality will have lower long-term fitness; thus, cyclical perturbations play an important role in driving the evolution of life history transition rates.

Period is not the only parameter of cycles, however. Particularly for seasons, cycle amplitude may also shape phenologies in important ways, and is shifting with climate change in many natural systems (e.g. seasonal CO_2_ cycle amplitude [71,72]). Amplitude of seasonal cycles may play two roles for evolution. First, amplitude is associated with intensity of disturbance, which can be explored with survivorship functions in my theoretical framework. If the pattern of stage-specific mortality associated with cyclical disturbance is clear, such as in *T. californicus* and many seasonal species, then heightened intensity of cyclical disturbance will likely increase strength of selection. Second, amplitude reflects the rate of environmental change within cycle phases. Rate of change may be important for cue-detection and plastic responses. For example many plants in seasonal environments are well known for tracking growing degree-days as a way of taking cues on the passing of the seasons [73]. In my theoretical framework, cyclical disturbances arrive without warning and simply incur repeated penalties on individuals and cohorts that had non-optimal life history strategies for the given regime. In reality there may be a number of continuously changing environmental variables in *T. californicus* pools such as salinity, and I cannot exclude the possibility that, like plants, birds, or many aquatic invertebrates, *T. californicus* possess biological mechanisms to use cues from continuously changing parameters to plastically alter their phenotypes. Nonetheless, I was able to predict variation in *T. californicus* life histories across a periodicity gradient in the environment without accounting for plasticity, suggesting that plastic responses might not have a strong effect on life history evolution in response to cyclicality. Future phenological work should directly compare the relative roles of demographic influences such as those discussed here and plastic response to cues that can be tracked along continuous cycles.

When considering phenological evolution in cyclical environments, the relative scaling of life cycles and environmental cycles becomes important. For instance, a perennial species must endure multiple seasonal cycle periods per generation. An annual species’ generation on the other hand fits within a single cycle period. In both cases, consequences of fitness-related phenotypes in one generation carry over to subsequent generations via intergenerational trade-offs in life histories [36], but the trajectory of evolution may differ between the two because of the number of cycle periods a generation experiences. Further, the model framework presented here assumes overlapping generations but many annual organisms have non-overlapping generations and synchronous phenologies. The evolutionary consequences of non-overlapping generations and synchronization in a population in cyclical environments should be explored further.

Phenology is the study of how life cycle schedules are fit to environmental cycles. A phenological trait is a manifestation of the aggregate life history strategy of a species [16], and expression timings of traits are ultimately controlled by transition rates between life history stages [20]. Phenological studies typically measure one representative phenotype such as flowering time in association with proximate drivers such as temperature or precipitation. But phenotypes covary and therefore one must consider trade-offs and competing selective forces with a whole-life perspective in order to understand the evolution of cyclical phenological traits. Here I placed such connections in the general context of environmental cycles, of which the annual seasonal cycle is one example, and tested mechanistic predictions on the relatively short timescale of tide cycles which yielded large amounts of data across many cycle periods and generations quickly. This framework provides a basis for analyzing, comparing, and predicting adaptive phenological shifts in changing seasonal environments.

## ACKNOWLEDGEMENTS

I thank G. Dwyer, M. Pascual, C.A. Pfister, T.D. Price, S.C. Stearns, J.T. Wootton, and two anonymous reviewers for helpful discussions and comments on earlier drafts. K. Miranda and T. Bowyer provided assistance in the field and with trait measurements. I am grateful to the Makah Tribe in Neah Bay, Friday Harbor Laboratories, and M. and D. Hurd for granting access to their lands for field data collection. This work was supported by University of Chicago Hinds Fund, DOE GAANN, and NSF LTREB grant (DEB-1556874).

## REFERENCES

1. Lampert W. 1989 The Adaptive Significance of Diel Vertical Migration of Zooplankton. Funct. Ecol. 3, 21–27. (doi:10.2307/2389671)

2. Sponaugle S, Cowen RK. 1994 Larval durations and recruitment patterns of two Caribbean gobies (Gobiidae): contrasting early life histories in demersal spawners. Mar. Biol. 120, 133–143. (doi:10.1007/BF00381949)

3. Post E, Stenseth NC. 1999 Climatic variability, plant phenology, and northern ungulates. Ecology 80, 1322–1339.

4. Keeley JE, Bond WJ. 1999 Mast Flowering and Semelparity in Bamboos: The Bamboo Fire Cycle Hypothesis. Am. Nat. 154, 383–391. (doi:10.1086/303243)

5. Schauber EM et al. 2002 Masting by eighteen New Zealand plant species: the role of temperature as a synchronizing cue. Ecology 83, 1214–1225.

6. Lytle DA, Poff NL. 2004 Adaptation to natural flow regimes. Trends Ecol. Evol. 19, 94–100. (doi:10.1016/j.tree.2003.10.002)

7. Post E, Forchhammer MC, Stenseth NC, Callaghan TV. 2001 The timing of life–history events in a changing climate. Proc. R. Soc. Lond. B Biol. Sci. 268, 15–23. (doi:10.1098/rspb.2000.1324)

8. Walther G-R, Post E, Convey P, Menzel A, Parmesan C, Beebee TJC, Fromentin J-M, Hoegh-Guldberg O, Bairlein F. 2002 Ecological responses to recent climate change. Nature 416, 389–395. (doi:10.1038/416389a)

9. Parmesan C, Yohe G. 2003 A globally coherent fingerprint of climate change impacts across natural systems. Nature 421, 37–42. (doi:10.1038/nature01286)

10. Edwards M, Richardson AJ. 2004 Impact of climate change on marine pelagic phenology and trophic mismatch. Nature 430, 881–884. (doi:10.1038/nature02808)

11. Parmesan C. 2006 Ecological and Evolutionary Responses to Recent Climate Change. Annu. Rev. Ecol. Evol. Syst. 37, 637–669.

12. Bradshaw WE, Holzapfel CM. 2008 Genetic response to rapid climate change: it’s seasonal timing that matters. Mol. Ecol. 17, 157–166.

13. Cleland EE, Chuine I, Menzel A, Mooney HA, Schwartz MD. 2007 Shifting plant phenology in response to global change. Trends Ecol. Evol. 22, 357–365. (doi:10.1016/j.tree.2007.04.003)

14. Thackeray SJ et al. 2010 Trophic level asynchrony in rates of phenological change for marine, freshwater and terrestrial environments. Glob. Change Biol. 16, 3304–3313. (doi:10.1111/j.1365- 2486.2010.02165.x)

15. Cohen JM, Lajeunesse MJ, Rohr JR. 2018 A global synthesis of animal phenological responses to climate change. Nat. Clim. Change 8, 224–228. (doi:10.1038/s41558-018-0067-3)

16. Post ES, Pedersen C, Wilmers CC, Forchhammer MC. 2008 Phenological sequences reveal aggregate life history response to climatic warming. Ecology 89, 363–370.

17. Both C, Van Asch M, Bijlsma RG, Van Den Burg AB, Visser ME. 2009 Climate change and unequal phenological changes across four trophic levels: constraints or adaptations? J. Anim. Ecol. 78, 73–83.

18. Chuine I. 2010 Why does phenology drive species distribution? Philos. Trans. R. Soc. Lond. B Biol. Sci. 365, 3149–3160. (doi:10.1098/rstb.2010.0142)

19. Richardson AD et al. 2010 Influence of spring and autumn phenological transitions on forest ecosystem productivity. Philos. Trans. R. Soc. Lond. B Biol. Sci. 365, 3227–3246. (doi:10.1098/rstb.2010.0102)

20. Forrest J, Miller-Rushing AJ. 2010 Toward a synthetic understanding of the role of phenology in ecology and evolution. Philos. Trans. R. Soc. Lond. B Biol. Sci. 365, 3101–3112. (doi:10.1098/rstb.2010.0145)

21. Visser ME, Caro SP, Oers K van, Schaper SV, Helm B. 2010 Phenology, seasonal timing and circannual rhythms: towards a unified framework. Philos. Trans. R. Soc. B Biol. Sci. 365, 3113–3127. (doi:10.1098/rstb.2010.0111)

22. Lande R, Engen S, Sæther B-E. 2017 Evolution of stochastic demography with life history tradeoffs in density-dependent age-structured populations. Proc. Natl. Acad. Sci. 114, 11582–11590. (doi:10.1073/pnas.1710679114)

23. Cook RE. 1979 Patterns of Juvenile Mortality and Recruitment in Plants. In Topics in Plant Population Biology, pp. 207–231. Palgrave, London.

24. Charlesworth B. 1994 Evolution in age-structured populations. Cambridge University Press Cambridge.

25. Caswell H. 2001 Matrix population models: construction, analysis, and interpretation. Sunderland, Mass.: Sinauer Associates.

26. Tuljapurkar S. 1985 Population dynamics in variable environments. VI. Cyclical environments. Theor. Popul. Biol. 28, 1–17. (doi:10.1016/0040-5809(85)90019-X)

27. Orzack SH. 1993 Life History Evolution and Population Dynamics in Variable Environments: Some Insights from Stochastic Demography. In Adaptation in Stochastic Environments, pp. 63–104. Springer, Berlin, Heidelberg.

28. Caswell H, Trevisan MC. 1994 Sensitivity Analysis of Periodic Matrix Models. Ecology 75, 1299–1303. (doi:10.2307/1937455)

29. Lytle DA. 2001 Disturbance Regimes and Life-History Evolution. Am. Nat. 157, 525–536. (doi:10.1086/319930)

30. Tuljapurkar S, Gaillard J-M, Coulson T. 2009 From stochastic environments to life histories and back. Philos. Trans. R. Soc. B Biol. Sci. 364, 1499–1509. (doi:10.1098/rstb.2009.0021)

31. Koons DN, Pavard S, Baudisch A, Metcalf JE. 2009 Is life-history buffering or lability adaptive in stochastic environments? Oikos 118, 972–980.

32. Gadgil M, Bossert WH. 1970 Life historical consequences of natural selection. Am. Nat. 104, 1–24.

33. Charnov EL, Schaffer WM. 1973 Life-history consequences of natural selection: Cole’s result revisited. Am. Nat. 107, 791–793.

34. Law R. 1979 Optimal life histories under age-specific predation. Am. Nat. 114, 399–417.

35. Michod RE. 1979 Evolution of life histories in response to age-specific mortality factors. Am. Nat. 113, 531–550.

36. Stearns SC. 1992 The Evolution of Life Histories. Oxford: Oxford University Press.

37. Roff DA. 1984 The Evolution of Life History Parameters in Teleosts. Can. J. Fish. Aquat. Sci. 41, 989–1000. (doi:10.1139/f84-114)

38. Hairston NG, Walton WE. 1986 Rapid evolution of a life history trait. Proc. Natl. Acad. Sci. 83, 4831–4833.

39. Reznick DN, Butler MJ, Rodd FH, Ross P. 1996 Life-history evolution in guppies (Poecilia reticulata) 6. Differential mortality as a mechanism for natural selection. Evolution 50, 1651–1660.

40. Ernande B, Dieckmann U, Heino M. 2004 Adaptive changes in harvested populations: plasticity and evolution of age and size at maturation. Proc. R. Soc. Lond. B Biol. Sci. 271, 415–423. (doi:10.1098/rspb.2003.2519)

41. Olsen EM, Heino M, Lilly GR, Morgan MJ, Brattey J, Ernande B, Dieckmann U. 2004 Maturation trends indicative of rapid evolution preceded the collapse of northern cod. Nature 428, 932–935. (doi:10.1038/nature02430)

42. Walsh MR, Post DM. 2011 Interpopulation variation in a fish predator drives evolutionary divergence in prey in lakes. Proc. R. Soc. Lond. B Biol. Sci. 278, 2628–2637. (doi:10.1098/rspb.2010.2634)

43. Orzack SH, Tuljapurkar S. 2001 Reproductive Effort in Variable Environments, or Environmental Variation Is for the Birds. Ecology 82, 2659–2665. (doi:10.1890/0012-9658(2001)082[2659:REIVEO]2.0.CO;2)

44. Metcalf CJE, Koons DN. 2007 Environmental uncertainty, autocorrelation and the evolution of survival. Proc. R. Soc. Lond. B Biol. Sci. 274, 2153–2160. (doi:10.1098/rspb.2007.0561)

45. Childs DZ, Metcalf CJE, Rees M. 2010 Evolutionary bet-hedging in the real world: empirical evidence and challenges revealed by plants. Proc. R. Soc. Lond. B Biol. Sci., rspb20100707. (doi:10.1098/rspb.2010.0707)

46. Fisher RA. 1930 The genetical theory of natural selection. Oxford, England: Clarendon Press.

47. Goodman D. 1982 Optimal Life Histories, Optimal Notation, and the Value of Reproductive Value. Am. Nat. 119, 803–823. (doi:10.1086/283956)

48. Koons DN, Grand JB, Zinner B, Rockwell RF. 2005 Transient population dynamics: relations to life history and initial population state. Ecol. Model. 185, 283–297.

49. Stott I, Townley S, Hodgson DJ. 2011 A framework for studying transient dynamics of population projection matrix models. Ecol. Lett. 14, 959–970.

50. Skellam JG. 1967 Seasonal periodicity in theoretical population ecology. In Proceedings of the 5th Berkeley symposium in mathematical statistics and probability, pp. 179–205.

51. Stearns SC. 1989 Trade-Offs in Life-History Evolution. Funct. Ecol. 3, 259–268. (doi:10.2307/2389364)

52. Pease CM, Bull JJ. 1988 A critique of methods for measuring life history trade-offs. J. Evol. Biol. 1, 293–303.

53. Powlik JJ. 1998 Seasonal abundance and population flux of Tigriopus californicus (Copepoda: Harpacticoida) in Barkley sound, British Columbia. J. Mar. Biol. Assoc. U. K. 78, 467–481.

54. Powlik JJ. 1999 Habitat characters of Tigriopus californicus (Copepoda: Harpacticoida), with notes on the dispersal of supralittoral fauna. J. Mar. Biol. Assoc. U. K. 79, 85–92.

55. Dethier MN. 1980 Tidepools as refuges: Predation and the limits of the harpacticoid copepod Tigriopus californicus (Baker). J. Exp. Mar. Biol. Ecol. 42, 99–111. (doi:10.1016/0022-0981(80)90169- 0)

56. Dybdahl MF. 1995 Selection on life-history traits across a wave exposure gradient in the tidepool copepod Tigriopus californicus (Baker). J. Exp. Mar. Biol. Ecol. 192, 195–210. (doi:10.1016/0022-0981(95)00063-W)

57. Roff DA, Heibo E, Vøllestad LA. 2006 The importance of growth and mortality costs in the evolution of the optimal life history. J. Evol. Biol. 19, 1920–1930.

58. Charnov EL, Turner TF, Winemiller KO. 2001 Reproductive constraints and the evolution of life histories with indeterminate growth. Proc. Natl. Acad. Sci. 98, 9460–9464. (doi:10.1073/pnas.161294498)

59. Schluter D, Price TD, Rowe L. 1991 Conflicting selection pressures and life history trade-offs. Proc R Soc Lond B 246, 11–17. (doi:10.1098/rspb.1991.0118)

60. Willett CS. 2010 Potential fitness trade-offs for thermal tolerance in the intertidal copepod Tigriopus californicus. Evolution 64, 2521–2534.

61. Kelly MW, Grosberg RK, Sanford E. 2013 Trade-Offs, Geography, and Limits to Thermal Adaptation in a Tide Pool Copepod. Am. Nat. 181, 846–854. (doi:10.1086/670336)

62. Salguero-Gómez R, Jones OR, Jongejans E, Blomberg SP, Hodgson DJ, Mbeau-Ache C, Zuidema PA, Kroon H de, Buckley YM. 2016 Fast-slow continuum and reproductive strategies structure plant life-history variation worldwide. Proc. Natl. Acad. Sci. 113, 230–235. (doi:10.1073/pnas.1506215112)

63. Cohen AA, Isaksson C, Salguero-Gómez R. 2017 Co-existence of multiple trade-off currencies shapes evolutionary outcomes. PLOS ONE 12, e0189124. (doi:10.1371/journal.pone.0189124)

64. Tuljapurkar SD. 1982 Population dynamics in variable environments. III. Evolutionary dynamics of r-selection. Theor. Popul. Biol. 21, 141–165. (doi:10.1016/0040-5809(82)90010-7)

65. Charlesworth B. 1990 Optimization models, quantitative genetics, and mutation. Evolution 44, 520–538.

66. Burton RS, Feldman MW. 1981 Population Genetics of Tigriopus Californicus. II. Differentiation Among Neighboring Populations. Evolution 35, 1192–1205. (doi:10.2307/2408132)

67. Burton RS. 1987 Differentiation and integration of the genome in populations of the marine copepod Tigriopus californicus. Evolution 41, 504–513.

68. Burton RS. 1997 Genetic evidence for long term persistence of marine invertebrate populations in an ephemeral environment. Evolution 51, 993–998.

69. Rathcke B, Lacey EP. 1985 Phenological Patterns of Terrestrial Plants. Annu. Rev. Ecol. Syst. 16, 179–214.

70. Remmel T, Tammaru T, Maegi M. 2009 Seasonal mortality trends in tree-feeding insects: a field experiment. Ecol. Entomol. 34, 98–106.

71. Keeling CD, Chin JFS, Whorf TP. 1996 Increased activity of northern vegetation inferred from atmospheric CO_2_ measurements. Nature 382, 146–149. (doi:10.1038/382146a0)

72. Angert A, Biraud S, Bonfils C, Henning CC, Buermann W, Pinzon J, Tucker CJ, Fung I. 2005 Drier summers cancel out the CO_2_ uptake enhancement induced by warmer springs. Proc. Natl. Acad. Sci. 102, 10823–10827. (doi:10.1073/pnas.0501647102)

73. Wolkovich EM et al. 2012 Warming experiments underpredict plant phenological responses to climate change. Nature 485, 494–497. (doi:10.1038/nature11014)

