## Supplementary Information for "Cyclical environments drive variation in life history strategies: a general theory of cyclical phenology"

*Supporting Information*

1. Model Description

2-4

2. Data Collection & Analysis

5-7

3. Supplementary Figures

8-11

4. Supplementary Tables

12-13

5. SI References

14

**1. MODEL DESCRIPTION**

**1.1 Continuous stage-structured population dynamics.**

First I express the dynamics of Juveniles and Adults of a population in the *absence of disturbance* with a simple system of ordinary differential equations:

$$\frac{dJ}{dt} = -(\mu+d)J+fA$$

$$\frac{dA}{dt} = \mu J-\gamma A$$

(S1)

where $\mu$ is the rate at which juveniles mature into reproducing adults, *d* is intrinsic juvenile mortality rate, *f* is reproductive rate of adults, and $\gamma$ is intrinsic adult mortality rate. This system, in matrix form, can be expressed as:

$$\mathbf{M}=\left[ \begin{matrix} -(\mu+d) & f \\ \mu& -\gamma\end{matrix} \right]$$

(S2)

The vector of solutions $<J\left( t \right), A\left( t \right)>$ can be expressed as $C_{1}v_{\lambda_{1}}e^{\lambda_{1}t}+C_{2}v_{\lambda_{2}}e^{\lambda_{2}t}$, thus

$$\begin{matrix} J\left( t \right)=C_{1}v_{\left( 1 \right)1}e^{\lambda_{1}t}+C_{2}v_{\left( 2 \right)1}e^{\lambda_{2}t} \end{matrix}$$

$$\begin{matrix} A\left( t \right)=C_{1}v_{\left( 1 \right)2}e^{\lambda_{1}t}+C_{2}v_{\left( 2 \right)2}e^{\lambda_{2}t} \end{matrix}$$

(S3)

where $v_{(i)j}$ is the $j^{th}$ element of eigenvector *i*, associated with eigenvalue $\lambda_{i}$ of $\mathbf{M}$.

To obtain the full solution I must solve for $\lambda_{1,}\lambda_{2}$ and $v_{(i)j}$ of $\mathbf{M}$, as well as the constants $C_{1}$and$C_{2}$ in terms of the eigenvalues and eigenvectors.

Let the abundance in each stage at $t=0$ be:

$$J\left( 0 \right)= C_{1}v_{\left( 1 \right)1}+ C_{2}v_{\left( 2 \right)1}$$

$$A\left( 0 \right)= C_{1}v_{\left( 1 \right)2}+ C_{2}v_{\left( 2 \right)2}$$

(S4)

Rearranging, I find:

$$C_{1}= \frac{A\left( 0 \right)- C_{2}v_{\left( 2 \right)2}}{v_{\left( 1 \right)2}}$$

$$C_{2}= \frac{J\left( 0 \right)- C_{1}v_{\left( 1 \right)1}}{v_{\left( 2 \right)1}}$$

(S5)

and substituting $C_{2}$ into $C_{1}$ I find:

$$C_{1}= \frac{A\left( 0 \right)- \frac{J\left( 0 \right)- C_{1}v_{\left( 1 \right)1}}{v_{\left( 2 \right)1}}v_{\left( 2 \right)2}}{v_{\left( 1 \right)2}}=\frac{A\left( 0 \right)}{v_{\left( 1 \right)2}}- \frac{\left( J\left( 0 \right)- C_{1}v_{\left( 1 \right)1} \right)v_{\left( 2 \right)2}}{v_{\left( 2 \right)1}v_{\left( 1 \right)2}}$$

$$C_{1}\left[ 1- \frac{{v_{\left( 1 \right)1}v}_{\left( 2 \right)2}}{v_{\left( 2 \right)1}v_{\left( 1 \right)2}} \right]=\frac{A\left( 0 \right)}{v_{\left( 1 \right)2}}- J\left( 0 \right)\frac{v_{\left( 2 \right)2}}{v_{\left( 2 \right)1}v_{\left( 1 \right)2}}$$

$$C_{1}\left[ \frac{v_{\left( 2 \right)1}v_{\left( 1 \right)2}-{v_{\left( 1 \right)1}v}_{\left( 2 \right)2}}{v_{\left( 2 \right)1}v_{\left( 1 \right)2}} \right]=\frac{A\left( 0 \right)}{v_{\left( 1 \right)2}}- J\left( 0 \right)\frac{v_{\left( 2 \right)2}}{v_{\left( 2 \right)1}v_{\left( 1 \right)2}}$$

$$C_{1}=\frac{A\left( 0 \right)v_{\left( 2 \right)1}-J(0)v_{\left( 2 \right)2}}{v_{\left( 2 \right)1}v_{\left( 1 \right)2}- v_{\left( 2 \right)2}v_{\left( 1 \right)1}}$$

(S6)

And similarly, I solve for $C_{2}$:

$$C_{2}=\frac{J\left( 0 \right)v_{\left( 1 \right)2}-A(0)v_{\left( 1 \right)1}}{v_{\left( 2 \right)1}v_{\left( 1 \right)2}- v_{\left( 2 \right)2}v_{\left( 1 \right)1}}$$

(S7)

Finally, substituting (S6) and (S7) into (S3), I find the solution for abundances of Juveniles and Adults at time *t* in the absence of disturbance:

$$J\left( t \right)= \frac{A\left( 0 \right)v_{\left( 2 \right)1}-J(0)v_{\left( 2 \right)2}}{v_{\left( 2 \right)1}v_{\left( 1 \right)2}- v_{\left( 2 \right)2}v_{\left( 1 \right)1}}v_{\left( 1 \right)1}e^{\lambda_{1}t}+ \frac{J\left( 0 \right)v_{\left( 1 \right)2}-A(0)v_{\left( 1 \right)1}}{v_{\left( 2 \right)1}v_{\left( 1 \right)2}- v_{\left( 2 \right)2}v_{\left( 1 \right)1}}v_{\left( 2 \right)1}e^{\lambda_{2}t}$$

$$A\left( t \right)= \frac{A\left( 0 \right)v_{\left( 2 \right)1}-J(0)v_{\left( 2 \right)2}}{v_{\left( 2 \right)1}v_{\left( 1 \right)2}- v_{\left( 2 \right)2}v_{\left( 1 \right)1}}v_{\left( 1 \right)2}e^{\lambda_{1}t}+ \frac{J\left( 0 \right)v_{\left( 1 \right)2}-A(0)v_{\left( 1 \right)1}}{v_{\left( 2 \right)1}v_{\left( 1 \right)2}- v_{\left( 2 \right)2}v_{\left( 1 \right)1}}v_{\left( 2 \right)2}e^{\lambda_{2}t}$$

|  |  | (S8) |
| --- | --- | --- |

This is the solution for demographic dynamics *in the absence of disturbance*, i.e. dynamics *between disturbance events*. Now I incorporate periodic disturbance and then lastly calculate population fitness as a result of periodicity of disturbance.

Note that the sole purpose of expanding the solution into the form (S8) is to isolate *t*, which will be used to incorporate periodic disturbance.

### 1.2 Incorporating periodic disturbance.

Let $J_{w}(t)$ and $A_{w}(t)$ describe the changes in the two stages through time before the first disturbance, and $J_{w+1}(t)$ and $A_{w+1}(t)$ before the second disturbance, and so on. Let *T* = length of each phase between disturbance events. Therefore, $J_{w}(T)$ and $A_{w}(T)$ are abundances immediately before the first disturbance, and $J_{w+1}(0)$ and $A_{w+1}(0)$ immediately after the first disturbance, and so on. Let $S_{J}$ and $S_{A}$ describe *survival rates* of each stage when disturbance occurs, so that:

$$J_{w+1}\left( 0 \right)= S_{J}J_{w}\left( T \right)$$

$$A_{w+1}\left( 0 \right)= S_{A}A_{w}(T)$$

(S9)

Thus,

$$\begin{matrix} {\overset{⃗}{N}}_{w+1} & =\left[ \begin{matrix} S_{J}J_{w}(T) \\ S_{A}A_{w}(T) \end{matrix} \right]={\overset{⃗}{N}}_{w} \\ & =\left[ \begin{matrix} S_{J}[\frac{A_{w}(0)v_{(2)1}-J_{w}(0)v_{(2)2}}{v_{(2)1}v_{(1)2}-v_{(2)2}v_{(1)1}}v_{(1)1}e^{\lambda_{1}T}+\frac{J_{w}(0)v_{(1)2}-A_{w}(0)v_{(1)1}}{v_{(2)1}v_{(1)2}-v_{(2)2}v_{(1)1}}v_{(1)2}e^{\lambda_{2}T}] \\ S_{A}[\frac{A_{w}(0)v_{(2)1}-J_{w}(0)v_{(2)2}}{v_{(2)1}v_{(1)2}-v_{(2)2}v_{(1)1}}v_{(1)2}e^{\lambda_{1}T}+\frac{J_{w}(0)v_{(1)2}-A_{w}(0)v_{(1)1}}{v_{(2)1}v_{(1)2}-v_{(2)2}v_{(1)1}}v_{(2)2}e^{\lambda_{2}T}] \end{matrix} \right]=\mathbf{P}\left[ \begin{matrix} J_{w}(0) \\ A_{w}(0) \end{matrix} \right] \\ & =\left[ \begin{matrix} \frac{J_{w}(0)S_{J}[(v_{(1)2}e^{\lambda_{2}T}v_{(1)2}-v_{(1)1}e^{\lambda_{1}T}v_{(2)2})]}{v_{(2)1}v_{(1)2}-v_{(2)2}v_{(1)1}}+\frac{A_{w}(0)S_{J}[(v_{(1)1}e^{\lambda_{1}T}v_{(2)1}-v_{(1)2}e^{\lambda_{2}T}v_{(1)1})]}{v_{(2)1}v_{(1)2}-v_{(2)2}v_{(1)1}} \\ \frac{J_{w}(0)S_{A}[(v_{(2)2}e^{\lambda_{2}T}v_{(1)2}-v_{(1)2}e^{\lambda_{1}T}v_{(2)2})]}{v_{(2)1}v_{(1)2}-v_{(2)2}v_{(1)1}}+\frac{A_{w}(0)S_{A}[(v_{(1)2}e^{\lambda_{1}T}v_{(2)1}-v_{(2)2}e^{\lambda_{2}T}v_{(1)1})]}{v_{(2)1}v_{(1)2}-v_{(2)2}v_{(1)1}} \end{matrix} \right]=\mathbf{P}\left[ \begin{matrix} J_{w}(0) \\ A_{w}(0) \end{matrix} \right] \end{matrix}$$

$$\mathbf{P}\boldsymbol{=}\left[ \begin{matrix} \boldsymbol{S}_{\boldsymbol{J}}\frac{\boldsymbol{[(}\boldsymbol{v}_{\boldsymbol{(1)2}}\boldsymbol{e}^{\boldsymbol{\lambda}_{\boldsymbol{2}}\boldsymbol{T}}\boldsymbol{v}_{\boldsymbol{(1)2}}\boldsymbol{-}\boldsymbol{v}_{\boldsymbol{(1)1}}\boldsymbol{e}^{\boldsymbol{\lambda}_{\boldsymbol{1}}\boldsymbol{T}}\boldsymbol{v}_{\boldsymbol{(2)2}}\boldsymbol{)]}}{\boldsymbol{v}_{\boldsymbol{(2)1}}\boldsymbol{v}_{\boldsymbol{(1)2}}\boldsymbol{-}\boldsymbol{v}_{\boldsymbol{(2)2}}\boldsymbol{v}_{\boldsymbol{(1)1}}} & \boldsymbol{S}_{\boldsymbol{J}}\frac{\boldsymbol{[(}\boldsymbol{v}_{\boldsymbol{(1)1}}\boldsymbol{e}^{\boldsymbol{\lambda}_{\boldsymbol{1}}\boldsymbol{T}}\boldsymbol{v}_{\boldsymbol{(2)1}}\boldsymbol{-}\boldsymbol{v}_{\boldsymbol{(1)2}}\boldsymbol{e}^{\boldsymbol{\lambda}_{\boldsymbol{2}}\boldsymbol{T}}\boldsymbol{v}_{\boldsymbol{(1)1}}\boldsymbol{)]}}{\boldsymbol{v}_{\boldsymbol{(2)1}}\boldsymbol{v}_{\boldsymbol{(1)2}}\boldsymbol{-}\boldsymbol{v}_{\boldsymbol{(2)2}}\boldsymbol{v}_{\boldsymbol{(1)1}}} \\ \boldsymbol{S}_{\boldsymbol{A}}\frac{\boldsymbol{[(}\boldsymbol{v}_{\boldsymbol{(2)2}}\boldsymbol{e}^{\boldsymbol{\lambda}_{\boldsymbol{2}}\boldsymbol{T}}\boldsymbol{v}_{\boldsymbol{(1)2}}\boldsymbol{-}\boldsymbol{v}_{\boldsymbol{(1)2}}\boldsymbol{e}^{\boldsymbol{\lambda}_{\boldsymbol{1}}\boldsymbol{T}}\boldsymbol{v}_{\boldsymbol{(2)2}}\boldsymbol{)]}}{\boldsymbol{v}_{\boldsymbol{(2)1}}\boldsymbol{v}_{\boldsymbol{(1)2}}\boldsymbol{-}\boldsymbol{v}_{\boldsymbol{(2)2}}\boldsymbol{v}_{\boldsymbol{(1)1}}} & \boldsymbol{S}_{\boldsymbol{A}}\frac{\boldsymbol{[(}\boldsymbol{v}_{\boldsymbol{(1)2}}\boldsymbol{e}^{\boldsymbol{\lambda}_{\boldsymbol{1}}\boldsymbol{T}}\boldsymbol{v}_{\boldsymbol{(2)1}}\boldsymbol{-}\boldsymbol{v}_{\boldsymbol{(2)2}}\boldsymbol{e}^{\boldsymbol{\lambda}_{\boldsymbol{2}}\boldsymbol{T}}\boldsymbol{v}_{\boldsymbol{(1)1}}\boldsymbol{)]}}{\boldsymbol{v}_{\boldsymbol{(2)1}}\boldsymbol{v}_{\boldsymbol{(1)2}}\boldsymbol{-}\boldsymbol{v}_{\boldsymbol{(2)2}}\boldsymbol{v}_{\boldsymbol{(1)1}}} \end{matrix} \right]$$

(S10)

where $\mathbf{P}$ is the matrix that relates the initial abundances of the two stages (${\overset{⃗}{N}}_{w}$) to those after disturbance (${\overset{⃗}{N}}_{w+1}$), via the length of the undisturbed phase, i.e. periodicity of disturbance ($T$), and the survival rate of each stage when disturbance occurs ($S_{J}$ and $S_{A}$).

**1.3 Fitness of a periodically disturbed population.**

Iterative multiplications of **P** to the initial population structure will simulate dynamics of the structure over many periodic disturbance cycles. Since periodicity in this model is not stochastic, **P** is equivalent to periodic matrix product models (Skellam 1967; Caswell 2001) whose dominant eigenvalue gives the long-run growth rate of the population. And by extension the dominant eigenvalue of **P** is an appropriate measure of the population’s fitness in a given regime *T*. In order to construct the fitness landscape across values of a life history trait across periodicity (*T*) regimes, I pass a value of a life history trait through **M**, allowing other life history traits to vary according to trade-off assumptions among traits, and calculate the dominant eigenvalue of the final matrix **P**. Eigenvalues of **P** are scaled against the highest eigenvalue per *T* in order to create a gradient per *T*. The life history trait per *T* that confers the highest fitness value is the optimal life history trait.

**2. DATA COLLECTION & ANALYSIS**

**2.1 *Tigriopus californicus.***

*T. californicus* has been a model species for marine phylogeography and population genetics research particularly due to its extensive latitudinal range and surprisingly strong local adaptation patterns (Burton & Feldman 1981; Burton *et al.* 1979; Edmands 2001; Edmands & Harrison 2003). Dense populations of *T. californicus* form in small pools, typically reaching tens of thousands of individuals in less than 10L of water (Dethier 1980; Powlik 1998, 1999). High tide levels periodically reach heights at which *T. californicus* pools occur and deliver wave disturbance. When waves flush through pools, normally free-swimming *T. californicus* dive and cling to the rock to avoid being washed out (Dethier 1980). Yet many are dislodged down to the mid and lower intertidal zones where predators such as sculpin and anemones feed on them quickly (Dethier 1980). Barriers of predation between pools are hypothesized as a mechanism that restricts exchange between populations and enhances local patterns of genetic structure even among nearby pools (Dethier 1980; Dybdahl 1995). The egg-to-egg generation time of *T. californicus* is about three weeks (Burton *et al.* 1979), and high tide disturbance periodicity is on the order of days to weeks. Thus disturbance period is on a comparable time scale to generation time. The exact disturbance period for a given pool varies due to local tide patterns and height of pool on the rocky shore. Once juveniles mature into reproductive maturity, males and females form mating pairs, and each female is fertilized only once in her life and continuously produces successive clutches of juveniles until she dies (Burton 1985).

I sampled 19 populations for local disturbance regime and life history traits. Sample populations span two regions of the Strait of Juan de Fuca in northern Washington that experience very different high tide patterns, one near the mouth (‘Neah Bay’), and one further inside the Strait (‘Friday Harbor’). I chose the two regions based on the hypothesis that Neah Bay is subject to clearer differences in spring and neap tides, creating longer periods between tidal disturbances in general but also a broader spectrum of periods among pools depending on pool height on the shores, and that Friday Harbor experiences much more consistent and higher high tides, subjecting populations there to shorter disturbance period regimes in general. I specifically quantified local disturbance period for each population using high-resolution temperature time series data to verify this regional hypothesis.

In *T. californicus*, rate of maturation (inverse of age at maturity) and fecundity are known to trade off strongly (Dybdahl 1995; Edmands & Harrison 2003; Willett 2010; Hong & Shurin 2015). This trade-off is in fact widespread and known to covary and evolve rapidly in conjunction with one another in many plants and animals (Stearns & Koella 1986; Stearns 1989). I also found a strong negative correlation between the two traits across my 19 populations (Fig. 2). Intrinsic survival rate of juveniles and adults are also known to trade off with maturation rate and adult fecundity, respectively, in *T. californicus* because energetically expensive osmoregulation and intracellular concentration of amino acids required in the harsh environment of rocky shores divert resources from reproduction and somatic growth (Goolish & Burton 1989; Dybdahl 1995).

**2.2 Local disturbance periods.**

Regional historical tide data were collected from National Oceanic and Atmospheric Administration’s National Data Buoy Center. I adhered HOBO (Onset Co.) temperature loggers at the bottom of each pool that recorded temperature data at 5-minute intervals for up to 4 months. Oceanic water temperatures are typically significantly lower than rocky shore pool temperatures, so sudden drops in temperature were taken as signals of tidal wave disturbance. When high tide rises enough to reach a given pool, multiple waves create clusters of signals. Therefore I used a sliding window and identified negative derivatives in the timeseries that were below a significance threshold compared to their neighbors (7 standard deviations below mean of all derivatives within that day). This method deals well with stochasticity within days, as well as day-to-day variations caused by weather effects, and was ground-truthed by occasional visual checks of tidal disturbance. I then calculated mean period between days that contain signals of wave disturbance per pool.

**2.3 Disturbed demographic dynamics.**

In order to quantify the general impact that high tide wave disturbance has on *T. californicus* populations, I chose a subset of five of the sample pools that had not experienced tidal disturbance for at least 3 days, and measured population structures 5 hours before an impending high tide disturbance and 5 hours after. To obtain comprehensive measurements of population structure and size at a given time point, I siphoned out entire volumes of water from pools using incrementally small plastic pipettes while squirting out small crevices and sucking out *T. californicus* individuals. I then vigorously swirled whole volumes in a uniform container (5-gallon bucket), took 50mL subsamples, and counted number of individuals in juvenile (nauplii and copepodites) and adult (both sexes) stages immediately under a scope. This method proved to yield consistent measurements of population structure, and provides a rare opportunity for sampling uniformly from known whole populations, quickly and easily. After the pre-disturbance structure measurements, I immediately returned all individuals to their pools to reinstate the populations. I chose nearby pools to determine whether pools exhibit high exchange of individuals when disturbance occurs, or if they exhibit a consistent pattern of structural change even if nearby, in line with previous studies that showed that mortality by predation in lower intertidal zones limit exchanges between pools.

**2.4 Life history measurements.**

I collected a random subsample of 30 mating adult pairs from each of the 19 populations after siphoning out the entire population. I kept each pair in an individual 6.9ml well filled with filtered local seawater (50μm) and 0.2g/l *Spirulina* powder in a common temperature environment. Once the pair copulated and separated, I extracted the male leaving only the female. In order to reduce potential lingering effects of variability in environmental conditions among home source pools, I let all fertilized females produce one clutch of eggs in the common garden setting before starting to measure fecundity. At 12 hour increments I checked for the production of a clutch of juveniles for every mother, transferred the mother to a fresh well if she had produced a clutch, recorded interval between clutches, and repeated until up to 8 clutches were produced per female or she died. I then calculated mean clutch interval per female for each population. Clutch size is known to be highly variable among *T. californicus* individuals, as well as among clutches per individual (Kelly *et al.* 2013), and clutch interval may be a better indicator of rate of reproduction that respond to selection experiments. I measured clutch size of the second clutch for a subset of mothers across populations. Rate of reproduction (*f* in the model) of each mother is inverse of mean clutch interval * mean clutch size. To measure age at maturity, I isolated 20 juveniles from the second clutch produced from each mother and reared them in a 6.9mL well with the same conditions as above (Kelly *et al.* 2013). At 12 hour increments I checked for visible egg sacs in females in each cluster. Every time a gravid female appeared I extracted her from the well so that she would not be counted twice. I recorded time to maturity for 5 females in a clutch from one mother. Rate of maturation (*μ* in the model) is the inverse of mean age at maturity.

**2.5 Parameterization for model analysis.**

Using my own life history data and data from various other studies that reported *T. californicus* life history traits, I set broad constraints on parameters through which I explored model behavior. Literature sources include studies in nature across the geographical range of *T.californicus*, or experimental studies that measured life history traits in control treatments as basis for measuring effects of factors such as UV or pollutants (Burton *et al.* 1979; Dethier 1980; Willett 2010; Barnett & Kontogiannis 1975; Scott 1995). These constraints are: *µ*=[0.01,0.05], *d*=[0.01,0.4], *f*=[4,17], and *γ*=[0.01,0.2]. Rates are daily rates.

3. SUPPLEMENTARY FIGURES

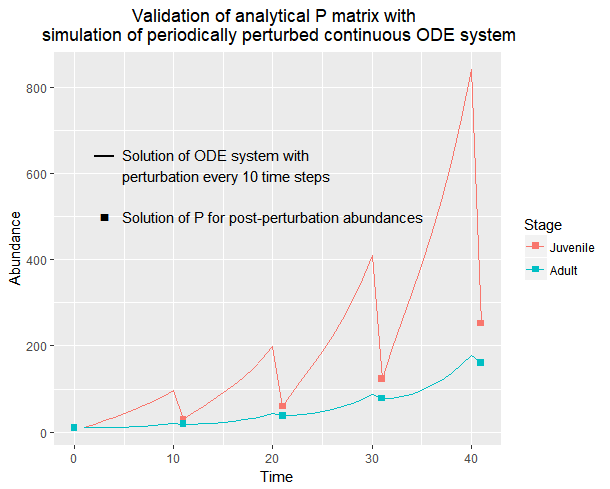

**Fig. S1.** Matrix multiplying an initial vector $\overset{⃗}{N}$ (whose elements represent stage-specific abundances) by **P** iteratively gives exactly the same solutions as simulating continuous stage-structured dynamics described by **M** for time *t* and then imposing stage-specific mortalities. This proof of concept shows that **P** is equivalent to periodic models which are typically comprised of discrete transition matrices that are multiplied. Here, I express dynamics between disturbances in continuous time, but stage transitions given after time *t* is equivalent to calibrating a discrete transition matrix for a projection interval *t*. Thus, time-invariant theoretical interpretations of matrix properties such as λ as long-run growth rate is justified.

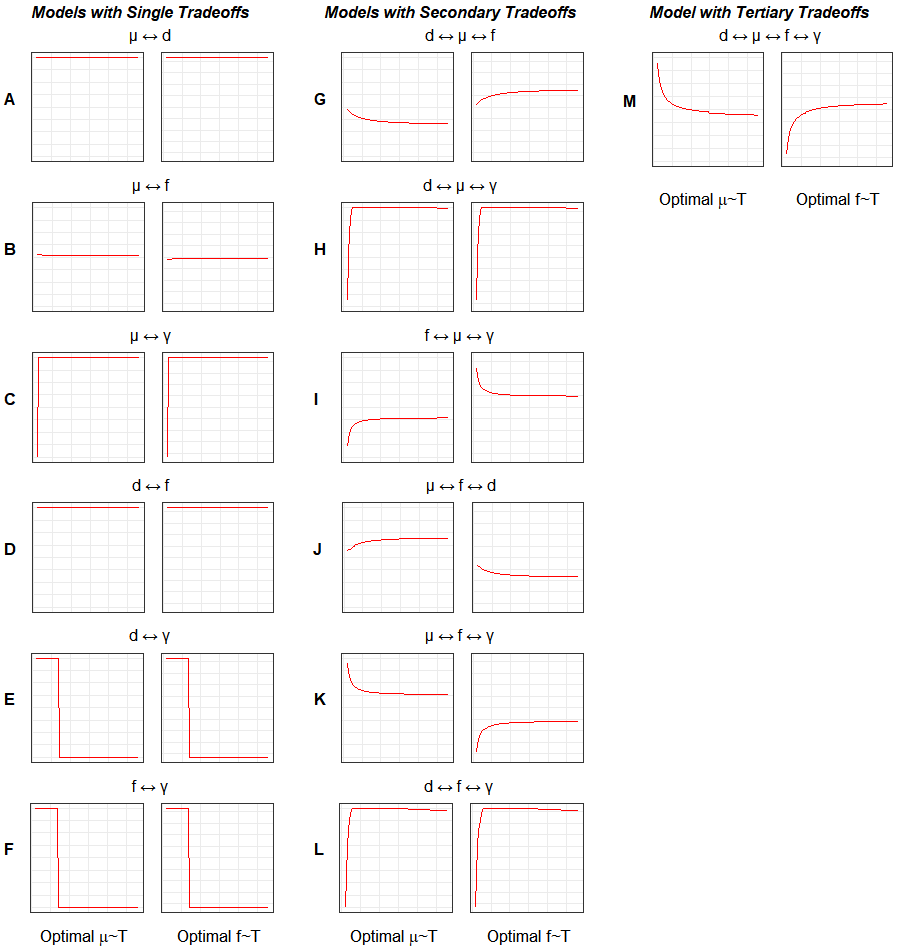

Fig. S2. Optimality curves of all model variants which include varying life history trade-off assumptions. Each model’s trade-off inclusion is denoted above the pair of plots by the double-sided arrow between traits. In each pair of plots the left panel is the curve representing optimal *μ* (fitness maximizing) across cycle period T, and right is optimal *f* across T. First column of paired plots contains models with single trade-offs, second column models with secondary trade-offs (but only ones that include *μ* or *f* out of all possible combinations among the four traits) and the last column the one model with tertiary trade-offs in which *μ* and *f* trade off with each other and with their respective stage mortalities *d* and *γ*. All realizations in this figure were run with S_A_ > S_J_ (0.9, 0.6), which represent stage-specific (Adults vs. Juveniles) mortalities associated with cyclical disturbance. Magnitude of S_A_ > S_J_ does not qualitatively change shapes of optimality curves.

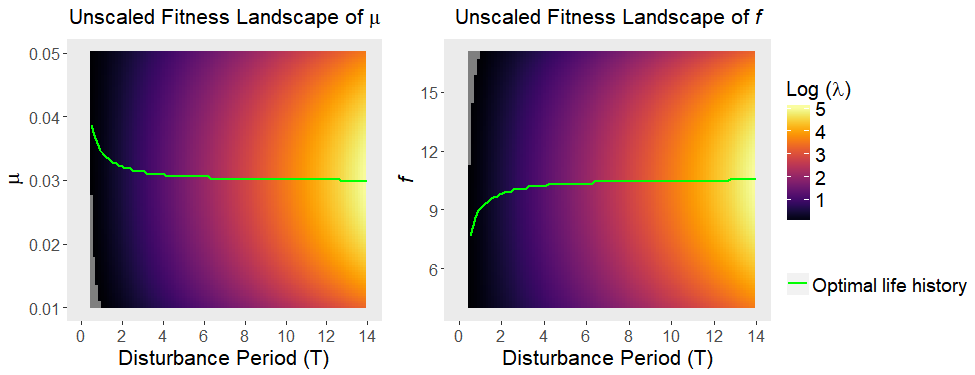

**Fig. S3.** Landscapes of log-transformed absolute fitness (eigenvalue of **P**) as opposed to relative fitness (normalized per T, as in Figure 2 in main text). Areas of log(λ) < 0 would represent negative long-term population growth, therefore potential troughs in evolutionary trajectories. These areas are shown in gray, appearing in the bottom-left corner of the *µ* landscape and top-left corner of the *f* landscape. Curves of optimal life history across T are identical to those in relative fitness landscapes, and do not cross areas of log(λ) < 0. The general incline in log(λ) as T increases is a result of higher long-term population growth due to less frequent disturbances.

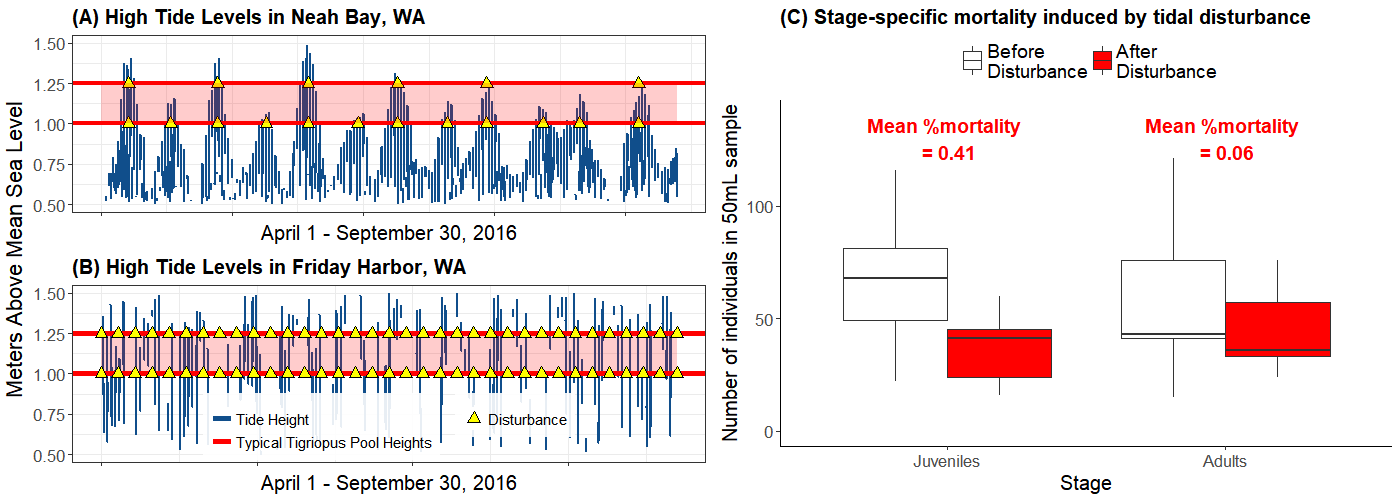

Fig. S4. High tide periodicity regimes in Neah Bay (A) and Friday Harbor (B) in northern Washington, which are near the mouth of and further inside the Strait of Juan de Fuca respectively. Data are hourly tide heights from NOAA’s National Data Buoy Center (Stations # 9449880 and # 9443090 respectively). Red lines that bound red shaded areas show typical range of heights where *T. californicus* populations are found on rocky shores. Yellow triangles show hypothetical disturbance events when high tide reaches height of a *T. californicus* pool, to illustrate that pools in Neah Bay are generally subject to longer periods and a wider spectrum of periods, and pools in Friday Harbor are generally subject to shorter periods. Disturbance events generally induce higher mortality in juveniles than adults (C), incurring not only population decline but also structural perturbation.

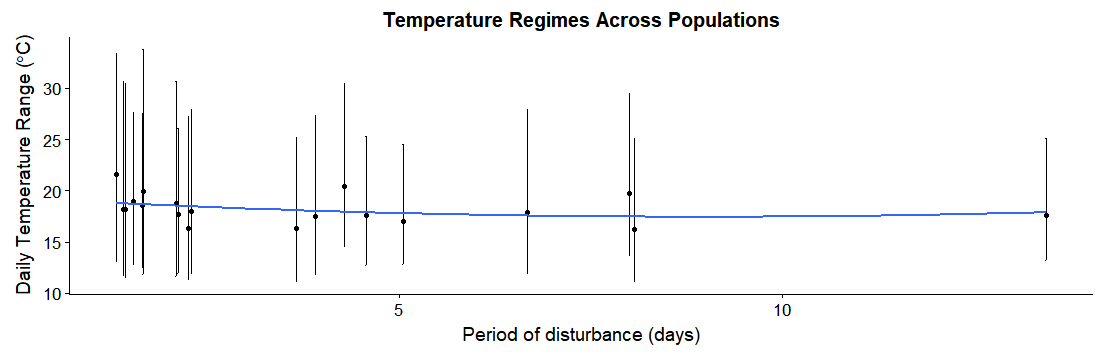

Fig. S5. Temperature regimes of sample populations plotted against pools’ tidal disturbance periods (data in Table S1), between June – September 2017. Point is mean daily mean temperature, and line spans mean daily minimum and mean daily maximum temperature per pool. Blue curve is second order polynomial fit on mean daily means. Generalized linear model (Mean daily mean ~ Period) showed non-significant difference across pools (*p = 0.28*).

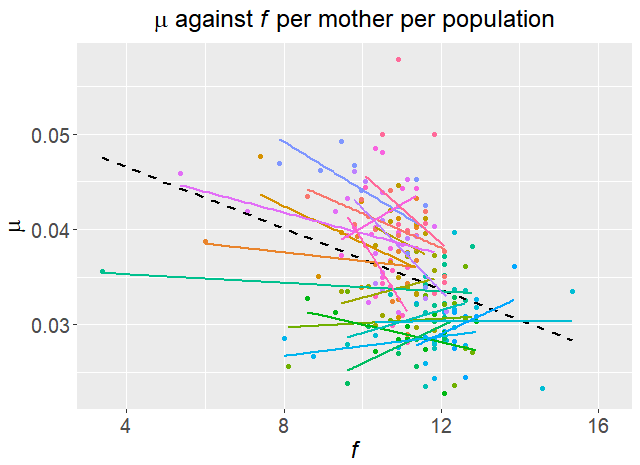

**Fig. S6.** Observed trade-offs between *µ* and *f* per mother in each population. Each point is a pair of a mother (used for *f* measurement) and her clutch (used for *µ* measurement), and 19 independent populations are represented by different colors. Dashed black line is the regression of *µ* against *f* across all populations.

4. SUPPLEMENTARY TABLES

Table S1. Temperature regimes of all 19 sample pools decomposed into mean daily min, mean daily mean, and mean daily max (plotted in Fig. S1). Population # is arbitrary codes of HOBO temperature loggers that were used to obscure association of each pool with its region or disturbance regime while conducting analyses. “FH” is Friday Harbor and “NB” is Neah Bay. Period is result of time series analysis for each pool’s 5-minute interval temperature data over the summer of 2017 that detected signals of wave disturbance and their intervals.

| **Population #** | **Region** | **Period** | **Daily Min °C** | **Daily Mean °C** | **Daily Max °C** |
| --- | --- | --- | --- | --- | --- |
| 1181396 | FH | 8 | 13.556 | 19.693 | 29.477 |
| 20148877 | FH | 1.646 | 11.875 | 19.961 | 33.808 |
| 20148879 | FH | 1.3 | 13.052 | 21.56 | 33.413 |
| 2276020 | FH | 1.64 | 12.38 | 18.575 | 27.585 |
| 2278765 | NB | 8.067 | 11.067 | 16.209 | 25.123 |
| 2292396 | NB | 3.903 | 11.88 | 17.513 | 27.349 |
| 2401944 | NB | 4.56 | 12.755 | 17.618 | 25.273 |
| 906044 | FH | 1.414 | 11.501 | 18.179 | 30.421 |
| 9742335 | NB | 13.444 | 13.238 | 17.588 | 25.143 |
| 20148874 | FH | 2.079 | 11.656 | 18.757 | 30.623 |
| 20148876 | FH | 1.39 | 11.642 | 18.201 | 30.701 |
| 20148878 | FH | 2.103 | 11.958 | 17.677 | 26.048 |
| 2276018 | FH | 1.519 | 12.706 | 18.947 | 27.657 |
| 2276206 | NB | 2.236 | 11.233 | 16.361 | 27.273 |
| 2278766 | NB | 6.667 | 11.838 | 17.898 | 27.939 |
| 2382989 | NB | 3.655 | 11.154 | 16.303 | 25.168 |
| 906035 | FH | 2.278 | 11.842 | 17.997 | 27.966 |
| 9742332 | NB | 5.043 | 12.827 | 16.996 | 24.568 |
| 9742339 | FH | 4.278 | 14.493 | 20.424 | 30.471 |

Table S2. Log-likelihoods of all models, obtained by maximum likelihood search through the space of S_A_ ≥ S_J_ (stage-specific mortalities associated with cyclical disturbance), while simultaneously fitting *μ* and *f*. Arrows denote trade-offs between life history traits. Model variants have different trade-off inclusions, but have the same number of estimated parameters because linear trade-offs were included computationally by setting parameter ranges of those traits in opposing (increasing vs. decreasing) order. Therefore model comparison criteria that penalize number of parameters were not needed. Highlighted (last model) is the best (likelihood-maximizing) model, which is reproduced as Figure 2 in the main text.

| Model (Trade-off) | Log-likelihood | Panel in Figure S2 |
| --- | --- | --- |
| *μ ↔ d* | *429.228* | *A* |
| *μ ↔ f* | *431.774* | *B* |
| *μ ↔ γ* | *429.228* | *C* |
| *d ↔ f* | *429.228* | *D* |
| *d ↔ γ* | *429.228* | *E* |
| *f ↔ γ* | *429.228* | *F* |
| *d ↔ μ ↔ f* | *441.527* | *G* |
| *d ↔ μ ↔ γ* | *431.431* | *H* |
| *f ↔ μ ↔ γ* | *429.228* | *I* |
| *μ ↔ f ↔ d* | *429.228* | *J* |
| *μ ↔ f ↔ γ* | *441.709* | *K* |
| *d ↔ f ↔ γ* | *441.120* | *L* |
| **d ↔ μ ↔ f ↔ γ** | **441.771** | ***M*** |

**5. SI REFERENCES**

Barnett, C.J. & Kontogiannis, J.E. (1975). The effect of crude oil fractions on the survival of a tidepool copepod, Tigriopus californicus. *Environ. Pollut. 1970*, 8, 45–54.

Burton, R.S. (1985). Mating system of the intertidal copepod Tigriopus californicus. *Mar. Biol.*, 86, 247–252.

Burton, R.S. & Feldman, M.W. (1981). Population Genetics of Tigriopus Californicus. II. Differentiation Among Neighboring Populations. *Evolution*, 35, 1192–1205.

Burton, R.S., Feldman, M.W. & Curtsinger, J.W. (1979). Population Genetics of Tigriopus californicus (Copepoda: Harpacticoida): I. Population Structure Along the Central California Coast. *Mar. Ecol. Prog. Ser.*, 1, 29–39.

Caswell, H. (2001). *Matrix population models: construction, analysis, and interpretation*. Sinauer Associates, Sunderland, Mass.

Dethier, M.N. (1980). Tidepools as refuges: Predation and the limits of the harpacticoid copepod Tigriopus californicus (Baker). *J. Exp. Mar. Biol. Ecol.*, 42, 99–111.

Dybdahl, M.F. (1995). Selection on life-history traits across a wave exposure gradient in the tidepool copepod Tigriopus californicus (Baker). *J. Exp. Mar. Biol. Ecol.*, 192, 195–210.

Edmands, S. (2001). Phylogeography of the intertidal copepod Tigriopus californicus reveals substantially reduced population differentiation at northern latitudes. *Mol. Ecol.*, 10, 1743–1750.

Edmands, S. & Harrison, J.S. (2003). Molecular and quantitative trait variation within and among populations of the intertidal copepod tigriopus californicus. *Evolution*, 57, 2277–2285.

Goolish, E.M. & Burton, R.S. (1989). Energetics of Osmoregulation in an Intertidal Copepod: Effects of Anoxia and lipid Reserves on the Pattern of Free Amino Accumulation. *Funct. Ecol.*, 3, 81–89.

Hong, B.C. & Shurin, J.B. (2015). Latitudinal variation in the response of tidepool copepods to mean and daily range in temperature. *Ecology*, 96, 2348–2359.

Kelly, M.W., Grosberg, R.K. & Sanford, E. (2013). Trade-Offs, Geography, and Limits to Thermal Adaptation in a Tide Pool Copepod. *Am. Nat.*, 181, 846–854.

Powlik, J.J. (1998). Seasonal abundance and population flux of Tigriopus californicus (Copepoda: Harpacticoida) in Barkley sound, British Columbia. *J. Mar. Biol. Assoc. U. K.*, 78, 467–481.

Powlik, J.J. (1999). Habitat characters of Tigriopus californicus (Copepoda: Harpacticoida), with notes on the dispersal of supralittoral fauna. *J. Mar. Biol. Assoc. U. K.*, 79, 85–92.

Scott, L.C. (1995). Survival and sex ratios of the intertidal copepod, Tigriopus californicus, following ultraviolet-B (290–320 nm) radiation exposure. *Mar. Biol.*, 123, 799–804.

Skellam, J.G. (1967). Seasonal periodicity in theoretical population ecology. In: *Proceedings of the 5th Berkeley symposium in mathematical statistics and probability*. pp. 179–205.

Stearns, S.C. (1989). Trade-Offs in Life-History Evolution. *Funct. Ecol.*, 3, 259–268.

Stearns, S.C. & Koella, J.C. (1986). The evolution of phenotypic plasticity in life-history traits: predictions of reaction norms for age and size at maturity. *Evolution*, 40, 893–913.

Willett, C.S. (2010). Potential fitness trade-offs for thermal tolerance in the intertidal copepod Tigriopus californicus. *Evolution*, 64, 2521–2534.
